# New Tools for Hop Cytogenomics: Identification of Tandem Repeat Families from Long-Read Sequences of *Humulus lupulus*

**DOI:** 10.1101/2020.02.03.931790

**Authors:** Katherine A. Easterling, Nicholi J. Pitra, Taylan B. Morcol, Jenna R. Aquino, Lauren G. Lopes, Kristin C. Bussey, Paul D. Matthews, Hank W. Bass

## Abstract

Hop (*Humulus lupulus* L.) is known for its use as a bittering agent in beer and has a rich history of cultivation, beginning in Europe and now spanning the globe. There are five wild varieties worldwide, which may have been introgressed with cultivated varieties. As a dioecious species, its obligate outcrossing, non-Mendelian inheritance, and genomic structural variability have confounded directed breeding efforts. Consequently, understanding genome evolution in Humulus represents a considerable challenge, requiring additional resources, including integrated genome maps. In order to facilitate cytogenetic investigations into the transmission genetics of hop, we report here the identification and characterization of 17 new and distinct tandem repeat sequence families. A tandem repeat discovery pipeline was developed using k-mer filtering and dot plot analysis of PacBio long-read sequences from the hop cultivar Apollo. We produced oligonucleotide FISH probes from conserved regions of HuluTR120 and HulTR225 and demonstrated their utility to stain meiotic chromosomes from wild hop, var. neomexicanus. The HuluTR225 FISH probe hybridized to several loci per nucleus and exhibited irregular, non-Mendelian transmission in male meiocytes of wild hop. Collectively, these tandem repeat sequence families not only represent unique and valuable new cytogenetic reagents but also have the capacity to inform genome assembly efforts and support comparative genomic analyses.

## INTRODUCTION

*Humulus lupulus* (hop) is a dioecious twining bine in the Cannabaceae family of flowering plants with a long history of cultivation (Neve, 1991; Moir, 2000) for various uses including medicine (as reviewed by Ososki and Kennelly, 2003; Bolton et al., 2019) and animal fodder (Siragusa et al., 2008), but is most commonly known as a flavoring agent in the brewing industry. The quest for complex taste and aromas in the rapidly expanding craft brewing industry has placed increasing demands on breeders to produce new varieties of plants with specific desirable traits as well as disease resistance (Kavalier et al., 2011; Easterling et al., 2018; Yan et al., 2019). However, hop presents multiple challenges to the production of new varieties due to its extended juvenile phase of two years to first flowers and its non-Mendelian inheritance patterns (Zhang et al., 2017).

Cytogenetic analysis of male meiosis in hop has revealed a tendency for unusual meiotic configurations such as multivalent chromosomal complexes (Sinotô, 1929; Winge, 1929; Shephard et al., 2000; Zhang et al., 2017). Recent work with 3D molecular cytology has shown that pervasive whole chromosome or segmental aneuploidy exists in hop and is exacerbated by passage through meiosis (Easterling et al., 2018). To date, there are limited cytological tools for assessing segregation patterns and establishing hop karyotypes (9 autosomes, XY). These tools have included telomere, 5S rDNA, HSR1 (Humulus subtelomeric repeat 1) (Karlov et al., 2003; Divashuk et al., 2011), and more recently HSR0 (Humulus subtelomeric repeat 0) (Easterling et al., 2018). Despite these advances, most genomes of model hop varieties remain to be sequenced, assembled, and fully annotated, except for unannotated, partial assemblies of Shinshu Wase, H lupulus var. cordifolius (Natsume et al., 2015) and Teamaker (Hill et al., 2017). Given the importance of cytogenetics in guiding studies of chromosomal structural genomics and the challenge presented by hop transmission genetics, more cytogenetic tools are needed.

Among the most useful tools for cytological chromosome analysis are FISH probes (Jiang, 2019). Of these, tandem repeat FISH probes are the most robust and have proven particularly useful for karyotyping and distinguishing chromosomes as illustrated for various eudicot and monocot plant species (Kato et al., 2004; Divashuk et al., 2011; Zuccolo et al., 2011; Weiss-Schneeweiss et al., 2015; Razumova et al., 2016; Novák et al., 2017; Easterling et al., 2018; Sun et al., 2018; Majtánová et al., 2019; Vondrak et al., 2019). Tandem repeat FISH probes specific for telomeres, centromeres, and knobs have also proven especially informative in 3D microscopic studies of meiosis (Bass et al., 1997; Birchler et al., 2007; Zuccolo et al., 2011; Howe et al., 2013; Higgins et al., 2014; Easterling et al., 2018; Majtánová et al., 2019).

Tandem repeat sequences clusters occur as tandemly repeated segments of DNA with characteristic unit repeat lengths. These clusters are among the fastest evolving components in genomes (Raskina et al., 2008; Weiss-Schneeweiss et al., 2015; Su et al., 2019) and are typically found in heterochromatic, noncoding DNA at centromeric, pericentromeric, or subtelomeric regions. Tandem repeats also have various and interesting chromosomal functions including transcriptional modulation of chromatin, centromere function, and meiotic drive (Garrido-Ramos, 2015; Dawe et al., 2018; Su et al., 2019). As such, they provide unique opportunities to study genome evolution and phylogenetic relationships (Dodsworth et al., 2015). Plants, particularly angiosperms, are characteristically rich in repetitive DNA, which can account for the vast majority of plant nuclear genomes (Mlinarec et al., 2019). Hop has been previously reported to contain around 34% repetitive elements in the assembled portions of the genome (Natsume et al., 2015), but that value will likely increase as more complete genome assemblies are produced.

Here we use long-read genomic sequences to find and characterize new families of hop tandem repeats as new tools for cytogenomic analysis of hop. We describe our discovery pipeline using k-mer filtering and dot plot analysis of single molecule long read sequence data from cultivar Apollo, resulting in identification of 17 new tandem repeat families. FISH probes from two of these, HuluTR120 and HuluTR225, are shown to detect loci in non-cultivated wild hops.

## METHODS

### Plant Materials, Collection, and Fixation

Male panicles were collected before pollen shedding and fixed in Farmer’s fluid as previously described (Zhang et al., 2017; Easterling et al., 2018). Wild hops, H. lupulus var. neomexicanus were collected from the Coronado National Forest in Arizona (U.S.A.). Plant SH2 was collected on Mt. Lemmon and plant TM2-82C was collected on Mt. Bigelow.

### Identification of Tandem Repeats in Long-Read PacBio sequences

Tandemly repeated sequences were discovered essentially using the approach previously described for the tandem repeat HSR0 (Easterling et al., 2018). Previously unreported details, parameters, and procedures are further described. DNA sequence input was hop (Apollo) genomic DNA from long-read PacBio DNA Single Molecule, Real-Time (SMRT) cells (libraries submitted Dec 2014, University of Washington PacBio Sequencing Services, Center https://pacbio.gs.washington.edu/) using single molecule sequencing without circular consensus error correction. The sequences from 32 SMRT cells had a library size range 3-20 kb, an average RQ (read quality) range of 81.5 - 82.55, and an Average Polymerase Mean Read Length (bp) ranged of 4,093 −5,048. For repeat detection, PacBio single molecule FASTA sequences greater than 5 kb (n=1,037,871) were subjected to k-mer analysis in which all 12mers were counted and sorted by abundance for each read. Sequences were filtered for retention if meeting the criterion where the fifth most repeated 12mer occurred at least eight times within a single read. This filter reduced the total list to 1,121 sequences, reflecting ~1000-fold enrichment. These k-mer filtered sequences were then used to produce a PDF document, referred to as the “HuluTR PDF book” in which each page shows the single-PacBio SMRT read dot plot for visual inspection along with a table of top 12mers using “ksift” (https://github.com/dvera/ksift) as previously reported (Easterling et al., 2018). In the resulting HuluTR PDF book (Supplementary Material 1), each page contains four items of information: (1) the page number, (2) the original FASTA file PacBio long-name sequence title, (3) the self-aligned YASS dot-plot, (4) the sequence and counts per read for the top 27 12mers sorted by abundance. The page numbers from the HuluTR PDF book have been used in our nomenclature as unique and single-number identifiers of the source read (-r#) sequences, replacing the long original FASTA titles. The 1,121 individual original format FASTA sequences (PacBio reads) are available in a supplemental file (Supplementary Material 2).

### Characterization of Tandem Repeat Families

Individual long reads that displayed prominent stripes in the dot plots were classified and placed into sequence similarity groups, also referred to as Tandem Repeat Families, using the online <u>YASS</u> dot-plot genome server (https://bioinfo.lifl.fr/yass/yass.php). Repeats were grouped into families if their pairwise dot-plots between two different reads returned “stripes” indicating repeating units of similar sequences between the two. For this, we used the default parameters from the YASS genome server which included Scoring matrix [match = +5, transversion = −4, transition = −3, other = −4 (composition bias correction)]; Gap costs [opening = −16, extension = −4]; [E-value threshold = 10]; [X-drop threshold = 30], and display DNA strain [fwd&rc] (Noe and Kucherov, 2005). To facilitate this process, we strung together sequences representing each TR family into a single “polySeq” FASTA file (Supplementary Material 3) and used it in each pairwise alignment with unclassified reads. New families (those not matching any of the repeats in the polySeq file) were added to the end of the polySeq file as they were discovered and included in blocks of sequence at 1kb intervals for ease of positional recognition in the YASS output dot plots. The Supplementary Material 3 file contains the full “polySeq34_v7” FASTA sequence with embedded locators, a table of synonyms to guide location to 1Kbp blocks, and individual YASS plots of the polySeq vs. each HuluTR consensus sequence. This family grouping process is illustrated in Figure 2.

For each TR family grouped by sequence similarity, we established an average consensus unit length based on the repeat spacing and results from the Tandem Repeats Finder server at https://tandem.bu.edu/trf/trf.html (Benson, 1999). Because of minor variation in the exact repeat lengths as determined by various methods, we rounded to the nearest 5bp and designated each HuluTR family accordingly, as summarized in Table 1. For each newly discovered HuluTR family, we obtained a single consensus sequence (Supplementary Material 4) from a representative read using the Tandem Repeats Finder online utility. Parameter settings used for TRF were default and as follows: alignment parameters (match = 2, mismatch = 2, indels = 7), minimum alignment score to report repeat = 50, Maximum period size = 1000, Maximum TR array size (bp, millions) = 2. The nomenclature used here denotes the TR family assignment with suffixes to specify single source reads as follows: “HuluTR120-r479” refers to <u>Hu</u>mulus <u>lu</u>pulus <u>T</u>andem <u>R</u>epeat of ~<u>120</u> bp PacBio read on page <u>479</u> of the HuluTR PDF book (Supplementary Material 2).

**Table 1.** Hop Tandem Repeats.

| HuluTR Family(A) | Repeat Size (bp)(B) | Parts per million (ppm)(C) | % A+T | Representative PacBio Read(D) | GenBank Accession |
| --- | --- | --- | --- | --- | --- |
| HuluTR385 (HSR1) | 385 | 232 | 61 | r55 | GU831574 |
| *HuluTR180 (HSR0) | 180 | 163 | 62 | r120 | MH188533.1 |
| *HuluTR120 | 120 | 69 | 67 | 782 | MN537570 |
| HuluTR335 (5SrDNA) | 335 | 63 | 55 | r243 | MN537579(E) |
| *HuluTR225 | 225 | 46 | 64 | r397 | MN537574 |
| *HuluTR060 | 60 | 34 | 59 | r91 | MN537567 |
| *HuluTR450 | 450 | 8 | 77 | r873 | MN537581 |
| *HuluTR135 | 135 | 5 | 42 | r253 | MN537571 |
| *HuluTR600 | 600 | 4 | 69 | r823 | MN537582 |
| *HuluTR390 | 390 | 2 | 79 | r15 | MN537580 |
| *HuluTR360 | 360 | 2 | 66 | r642 | MN537578 |
| *HuluTR240 | 240 | 2 | 71 | r1001 | MN537575 |
| *HuluTR185 | 185 | 2 | 49 | r424 | MN537573 |
| *HuluTR100 | 100 | 2 | 62 | r983 | MN537569 |
| *HuluTR350 | 350 | 1 | 67 | r625 | MN537577 |
| *HuluTR280 | 280 | 1 | 69 | r934 | MN537576 |
| *HuluTR150 | 150 | 1 | 60 | r390 | MN537572 |
| *HuluTR070 | 70 | 1 | 73 | r541 | MN537568 |
| *HuluTR055 | 55 | 1 | 64 | r292 | MN537566 |
| *HuluTR050 | 50 | 1 | 57 | r33 | MN537565 |

For analysis of monomer divergence within and between reads of HuluTR210, we extracted 11 monomers from an internal contiguous cluster for each of 10 reads. These were analyzed using a multiple sequence alignment tool, Clustal Omega (Clustal 2.1, https://www.ebi.ac.uk/Tools/msa/clustalo/). The resulting Percent Identity Matrix was imported into MS Excel and the sequence identity values were visualized for the individual monomers or their read-to-read averages (Fig. 3) using the Conditional Formatting tool with 2-Color Scale set from 40 (black) to 70 (yellow).

### FISH and 3D Cytology

Male meiocytes from hop plants were prepared, analyzed, and imaged using 3D deconvolution microscopy as previously described (Easterling et al., 2018). Nucleoli were measured using the Measure Distances program in the DeltaVision Software. Their diameter measurements were taken from central optical sections of each nucleolus, which are primarily spherical. Seventeen nucleoli were measured for cells with only one nucleolus (n=17 cells) and twelve were measured for cells with two nucleoli (n=8 cells). Average diameters were converted to volume in cubic microns.

Tandem repeat oligo names, sequences, and associated dyes utilized and reported in FISH experiments are as follows: “TR120Y” is 5’ -[ATTO647N]-GAGCACGAGATATTGATAAAAA, “TR225Y” is 5’-[ATTO647N]-TTAGTGCAATGTTATCTAGT. Additional sequences for oligo FISH probes and synthetic consensus sequences were designed and are listed here (Table 2) in order to provide new information as additional tools for hop cytogenetics. The synthetic consensus sequences were made (GenScript Biotech Corp.) and inserted into plasmids to enable their use as templates to make FISH probes via conventional labeling techniques. These plasmids (pHTR120syn, pHTR225syn, pHTR600syn, pHTR390syn, an pHTR060syn) and their descriptions are available from AddGene (addgene.org).

**Table 2.**
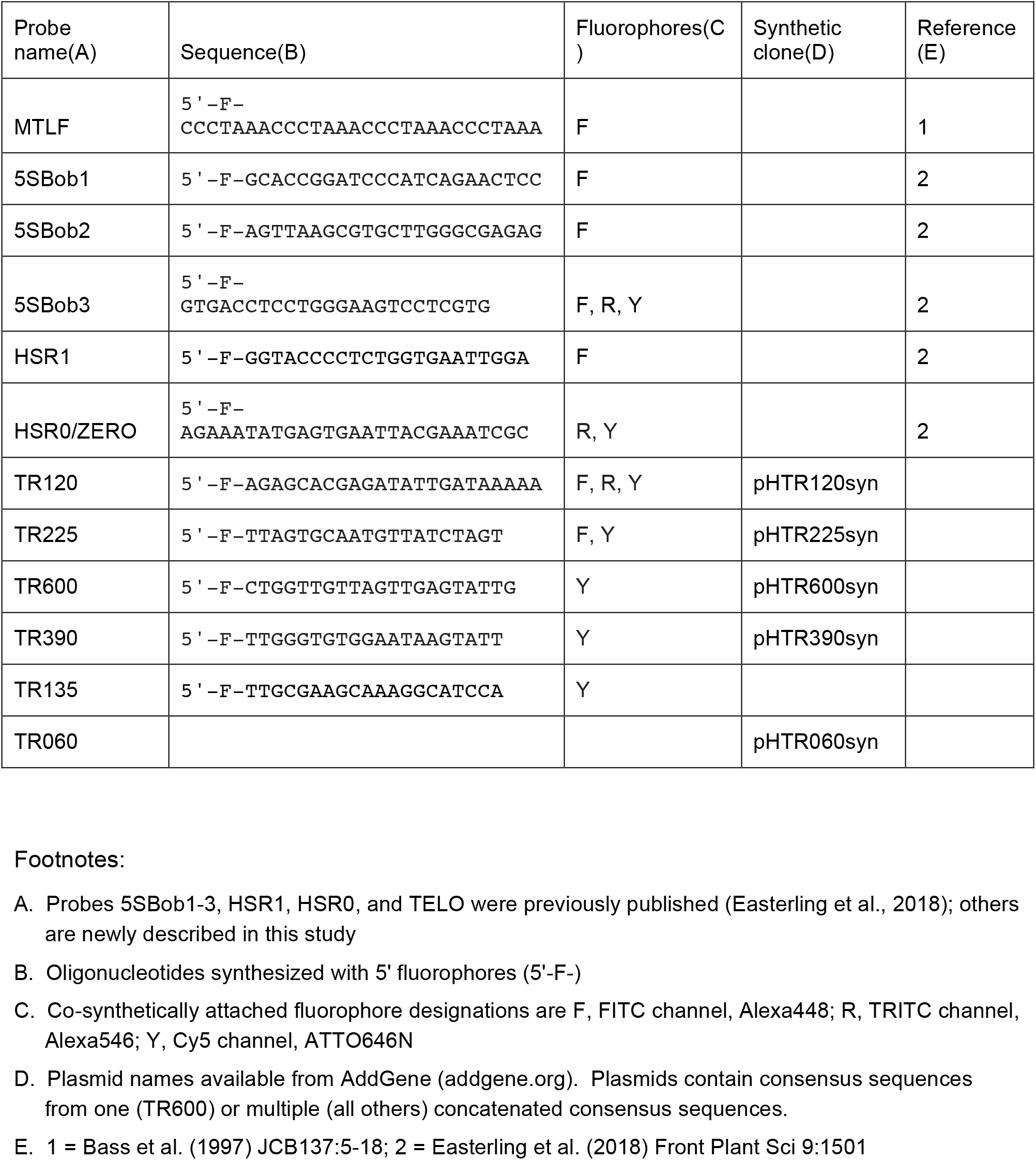
HuluTR Family-Specific Oligo FISH probes.

## RESULTS

In this study, we set out to develop new FISH probes that can be used for cytogenetic tracking of individual chromosomes in the Humulus lupulus species. To date, there exist only a few such probes including those for rDNA repeats and other tandemly repeated clusters. These have served to establish basic hop karyotypes, but more cytogenomic information is necessary in order to further delineate individual chromosomes and integrate physical and linkage maps for this group of plants.

### Finding Tandem Repeats with K-mer and Dot Plot analyses of PacBio Long-Read Sequence Data

We and others have successfully mined sequence data to identify tandem repeats that have been developed into FISH probes (Novak et al., 2013; Sevim et al., 2016; Novák et al., 2017; Easterling et al., 2018; Mlinarec et al., 2019). Here we carried out a thorough analysis of PacBio Single Molecule, Real-Time (SMRT) reads (n=1,037,871 reads), each consisting of sequences greater than 5,000 bp long. These reads, from 2014, produced single molecule DNA sequence, not circular consensus corrected, with an estimated error rate of ~10% based on alignments with a telomeric test case (Supplementary Material 5). We used k-mer analysis to screen for repetitive sequences. The criterion used was that the 5th most abundant 12 base k-mer for any single read be present eight or more times. This computational filter resulted in a list of 1,121 reads which were visualized as self-aligned Dot Plots using the YASS program (Noe and Kucherov, 2005) as summarized in Figure 1. Self-aligned dot plots display the same sequence on the X and Y axis and produce a diagonal line indicative of their sequence similarity, or complete identity in this case. Repetitive sequences within a read show up as off-diagonal stripes at a frequency and spacing that reflect their abundance and repeat unit size.

**Figure 1.**
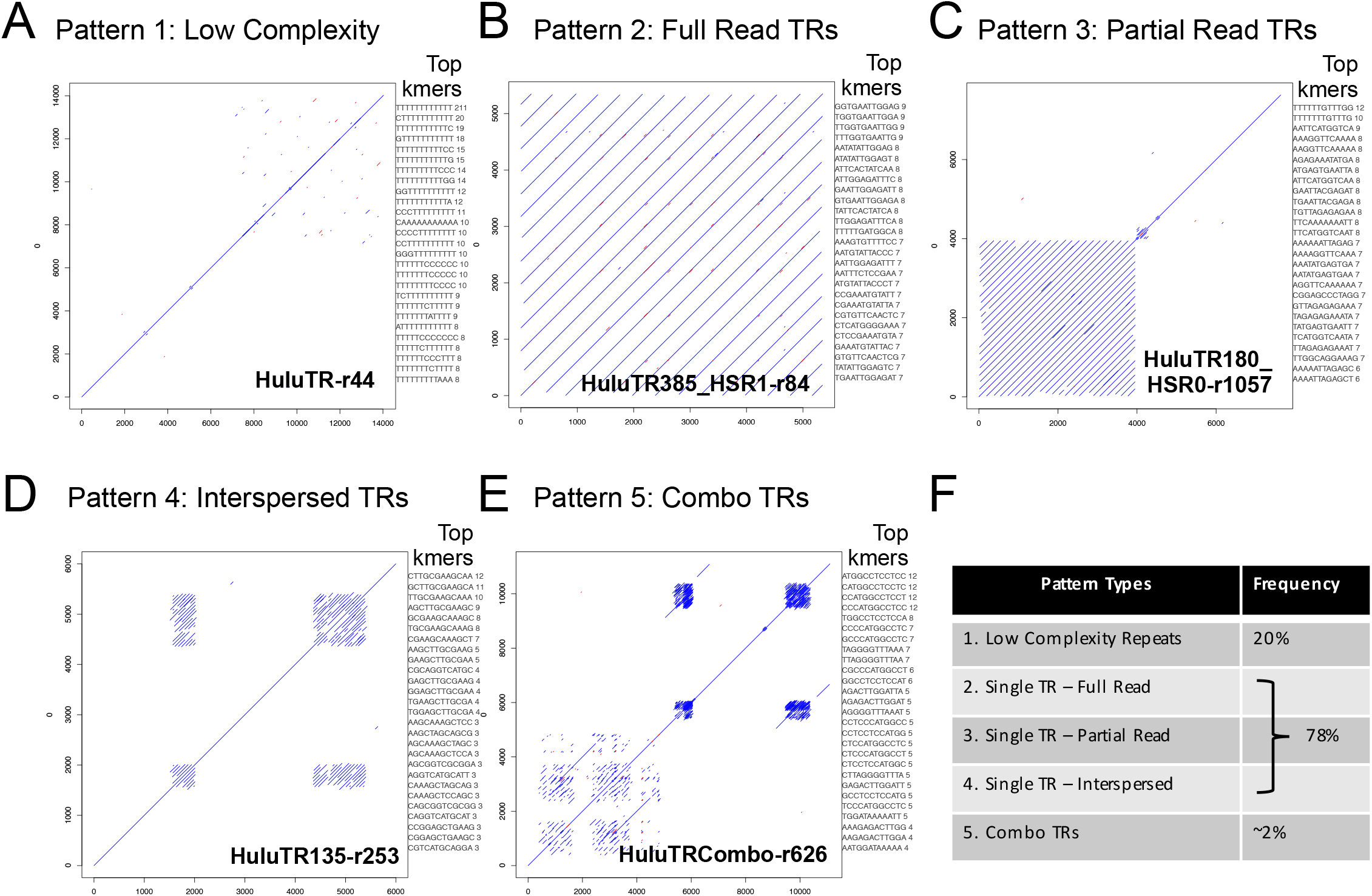
Dot plot outputs of k-mer analysis, showing different pattern types. PacBio Single Molecule, Real-Time (SMRT) DNA sequences were screened for tandem repeats. For each read, a self-aligned dot-plot is shown along with the top 20 12-mers for that read using k-mer analysis. The off-diagonal stripes represent internal tandem repeats. (A) Pattern 1: Example showing the HuluTR-r44 read showing no off-diagonal stripes, indicating lack of long or regular tandem repeats, referred to here as low-complexity repeats. (B) Pattern 2: Example showing the HuluTR385_HSR1-r84 read in which the tandem repeats occupy an entire read. (C) Pattern 3: Example showing the HuluTR180_HSR0-r1057 read in which the tandem repeats occupy part of the read. (D) Pattern 4: Example showing the HuluTR135-r253 read in which the tandem repeats occupy multiple but interspersed regions of the read. (E) Pattern 5: Example showing the HuluTRCombo-r626 read in which the more than one tandem repeat family is present in the same read. (F) Percentages of each of the five HuluTR pattern types from the 1,121 k-mer-filtered reads.

**Figure 2.**
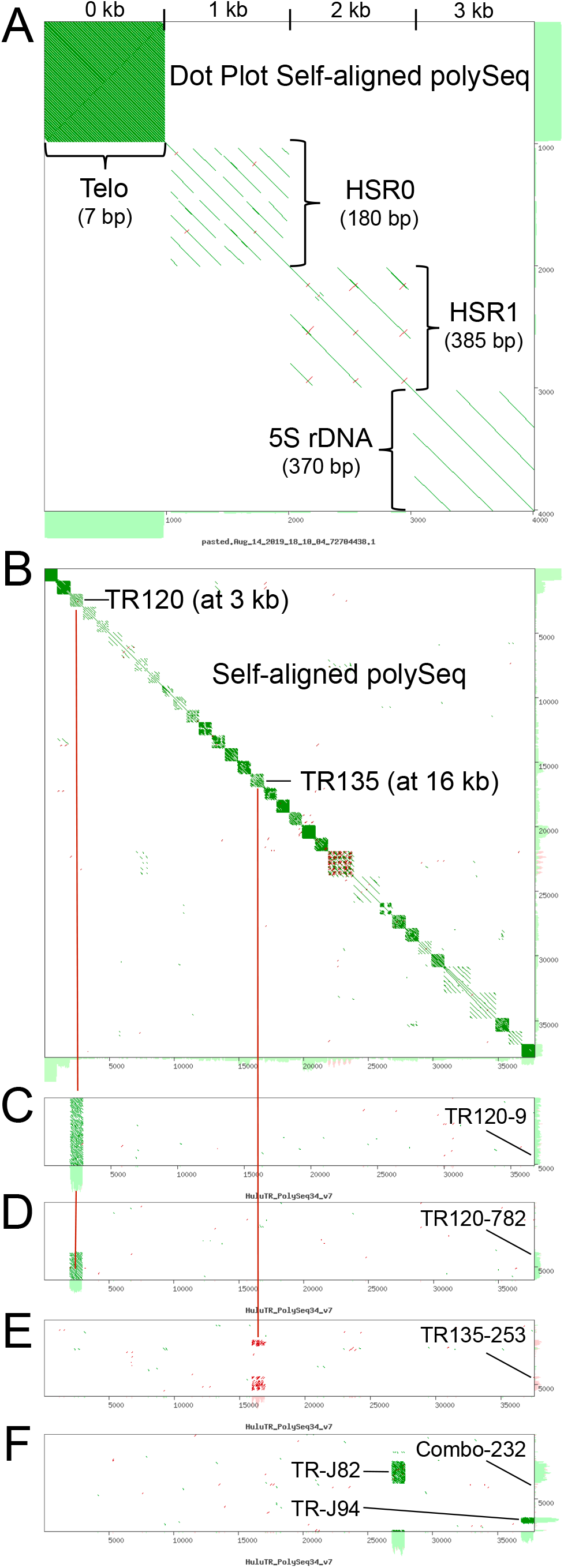
Using polySeq to define HuluTR families. A concatenation of consensus sequences for each known repeat family was made for use in the YASS dot-plot analysis to group tandem repeats into existing or new families, one read at a time. (A) Previously known repeats, Telo, HSR0, HSR1 and 5S rDNA showing 1 kb blocks in a dot plot of self-aligned polySeq. (B) Dot plot output of all 34 polySeq repeat sequences results in our study (FASTA sequence in Supplemental Material 3). Examples of read matching are indicated by the dark red bars denoting alignment to polySeq regions at 3 kb, corresponding to TR120, and at 16 kb, corresponding to TR 135. (C) HuluTR120-r9 with a Full read TR pattern, matching with the polySeq at 3 kb. (D) HuluTR120-r782 with a Partial read TR pattern, matching with the polySeq at 3 kb. (E) HuluTR135-r253 with an Interspersed TR pattern, matching with the polySeq at 16 kb. (F) HuluTRCombo-r232 with a Combo TR pattern, matching with the polySeq at 27 and37 kb.

**Figure 3.**
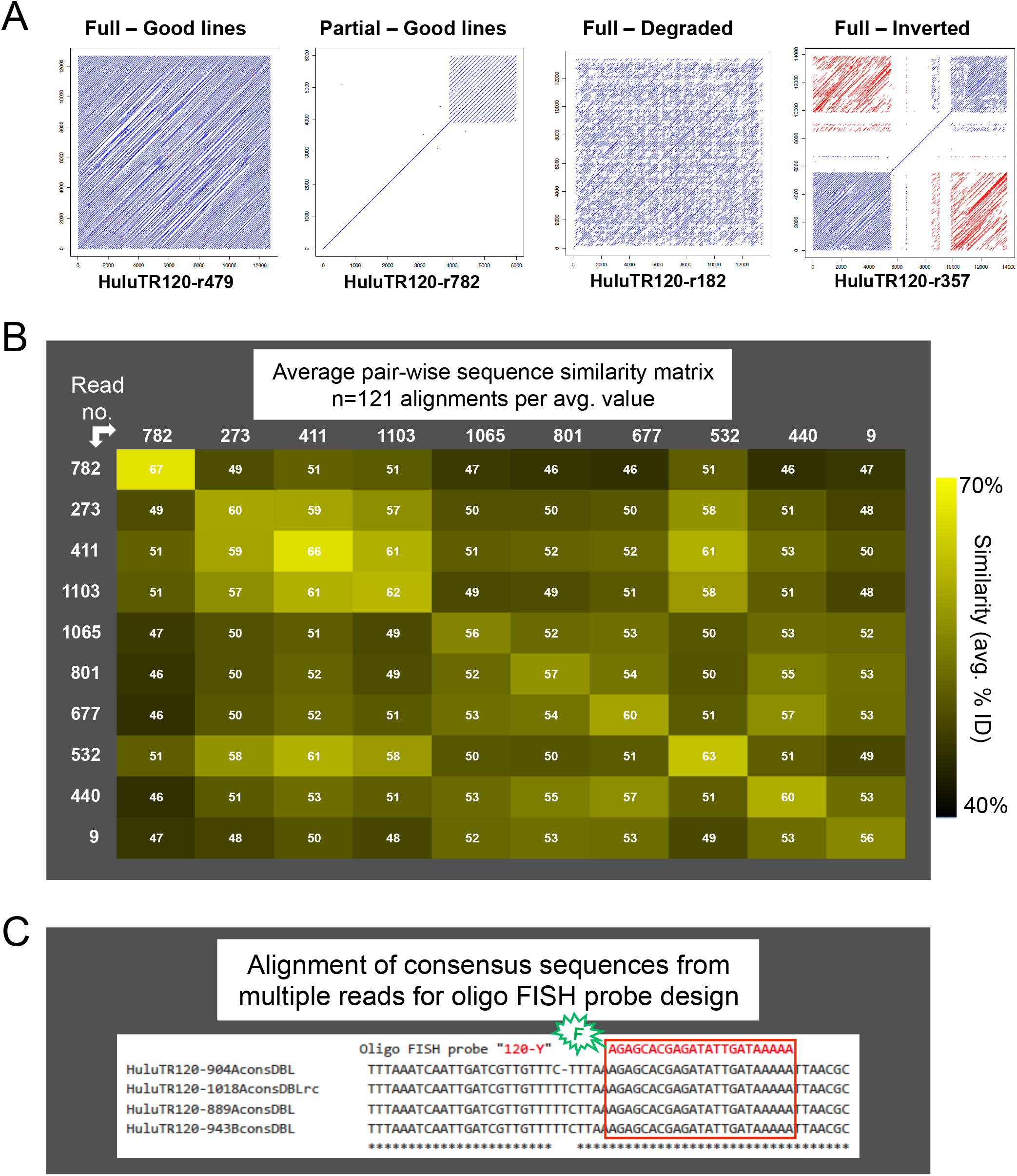
Sequence variation in the HuluTR120 family. (A) Differences observed in striping patterns from reads that match the HuluTR120 family. Stripe patterns are designated according to the aspects of the off-diagonal alignment stripes and designated as “Good” or “Degraded” in reads with Full or Partial TR occupancy. (B) Pair-wise heat map matrix of average sequence similarity (% identity) between all the monomers between any two reads where the diagonal boxes report within-read average. The numbers on the left and across the top identify the PacBio source read. The color scheme is shown to the right. (D) CLUSTAL Omega sequence alignment example for identification of short sequence OLIGO for fluorescent labeling and subsequent FISH probe design.

Several types of repeat sequence patterns were observed among the 1,121 reads that passed k-mer screen. The dot-plot pattern types can be grouped as those with low complexity and no obvious tandem repeats (Fig. 1A, no off-diagonal line/stripes) or those with clear tandem repeats, which present as stripes and fall into several subgroups (Fig. 1B-E). The spacing between the stripes resulting from tandem repeats is proportional to the repeat unit length, and these plots provide easy to interpret summary diagrams. Low complexity reads (e.g. Fig. 1A) comprised ~20% of the k-mer filtered reads, but were not further analyzed. They included homopolymeric runs of single or simple sequence repeats or microsatellites. In contrast, desirable reads of larger tandem repeats showed conspicuous dot-plot striping. These could be further subdivided into groups where the tandem repeats fill an entire read (Full Read TRs, Fig. 1B), a single portion of a read (Partial Read TRs, Fig. 1C), multiple but separate patches of the same repeat in a read (Interspersed TRs, Fig. 1D), or separate patches of dissimilar repeats in a read (Combo TRs, Fig. 1E). The reads with the Combo TRs account for ~2% of the full k-mer set (Fig. 1F) and often include repetitive sequence clusters with relatively short repeat lengths of ~30-50 bp. These Combo TR reads, like the low complexity reads were not prioritized for further analysis.

By mining long-read sequence data, our pipeline identified nearly 900 PacBio SMRT reads with tandem repeats with full, partial, or interspersed TR patterns. Among these were reads housing known tandem repeat families (5S rDNA, HSR1, HSR0) and those housing new uncharacterized tandem repeat families. To consolidate and sort out the newly discovered TR families, we propagated the page number from the 1,121-page PDF file, allowing for downstream grouping on the basis of sequence similarity.

### Defining HuluTRs: the Tandem Repeat Families of Hop

We systematically defined TR sequence families by grouping the tandem repeats according to sequence similarity criteria using dot plot analysis as summarized in Figure 2. The process is illustrated for four previously known tandem repeats (Fig. 2A): telomere, HSR0, HSR1, and 5S rDNA. For each TR, a 1kb block representing a TR family was made by a concatenation of a single repeating units or consensus sequence repeats. These 1-kb TR family-specific sequence blocks provide convenient visual delineations on the dot plot axes. These blocks help with assigning new repeat hits to matching families and were further concatenated to produce a file called “polySeq”. The resulting 4-TR polySeq (shown as self-aligned in Fig. 2A) was used as one of the two inputs to screen new reads by dot plotting, one at a time. For each new, uncharacterized read (those not matching sequences in the existing polySeq), we gave them a name (based on unit repeat length or discovery number) and included them in the polySeq file as a 1kb block of repeats, or 2kb blocks for large repeats. This process was repeated for each read, eventually producing a polySeq set of 34 distinct TR families (Supplementary Material 2), shown as a self-aligned dot plot (Fig. 2B). The TR-family assignment procedure is illustrated for four different reads in panels C-F (Fig. 2). For these examples, the dot plot shows the result with the polySeq on the X-axis and the query read on the Y-axis. The TR patterns shown include examples designated full read TRs (Fig. 2C), partial read TRs (Fig. 2D), interspersed TRs (Fig. 2E), or combo TRs (Fig. 2F). The fact that the self-aligned polySeq-34 dot plot (Fig. 2, panel B) as well as the pairwise queries (Fig. 2C-F) show sequence similarity striping within but not between the different TR families demonstrates the specificity of this approach, even when using single molecule reads with relatively high intrinsic error rates.

From the 34 hop TR repeat families examined, a total of 18 new HuluTR families were reported, choosing those with larger repeats (> 50 bp) and tendency to occur as the only TR cluster in a given read (e.g. reads in Fig 2 panels C-E, but not F). Tandem repeat families are named and listed in Table 1, sorted sequentially by relative abundance then by repeat length. For read-specific nomenclature, “HuluTR120-r9” refers to “<u>Hu</u>mulus <u>lu</u>pulus <u>T</u>andem <u>R</u>epeat family ~<u>120</u> bp from the PacBio sequence <u>r</u>ead on page <u>9</u> of the HuluTR PDF book (Supplementary Material 1). The six most abundant TR families found in the library range from 34 to 232 parts per million, and included previously known sequences HSR1, HSR0, and 5S rDNA, and newly discovered sequences, HuluTR120, HuluTR225, and HuluTR060. Their relative abundance makes them good candidates for FISH probes. The other families were found to occur at a lesser frequency, including six that were found in only one read.

Several of these TR clusters feature a high %A+T (AT content), as is often observed for tandemly-repeated macrosatellite sequences (Garrido-Ramos, 2017). The average AT content for the 18 sequence families listed ranged from an unusually low value of 42% for HuluTR135 to a high value of 79% for HuluTR390. The mean and median AT content for these sequence families is ~64%, higher than the 59.2% AT content calculated for all of the sequences in the PacBio Apollo DNA sequence dataset. This TR discovery approach expands the number of characterized hop TR sequence families by 10-fold and this methodology could be readily applied to other plant species for which long-read sequence datasets are available.

### Development of New Tandem Repeat FISH probes: Selection of Representative Sequences for TR FISH probe production

Once the tandemly repeated DNA sequences were categorized by family, we aimed to produce representative oligonucleotide FISH probes for cytogenetic detection of the corresponding chromosomal loci. Oligo FISH probes are advantageous because of their small size, uniformity of labeling, and consistency across experiments. The goal of identifying the best region of a tandem repeat family to use as a FISH probe is complicated by considerable sequence variation that is commonly observed in tandem repeat sequence families (Dennis and Peacock, 1984). For instance, as summarized in Figure 3 for sequences of HuluTR120 family, we observed variation from one read to another in the dot plot striping patterns. We consider continuous, parallel stripes to reflect tandem repeats with a high degree of similarity (Fig. 3A, 1st two plots). Such sequences were given high priority for probe development. However, some reads exhibited a more degraded appearance of stripes (Fig. 3, 3rd plot), which we interpret as having undergone sequence divergence, and were excluded from use in probe development.

To illustrate the range of sequence similarity variation both between and within reads, we selected 10 reads assigned to the HuluTR120 family (Fig. 3B). For each read, we extracted an internal, contiguous 11-repeat block of HuluTR120 monomers and separated them to quantify all possible monomer-to-monomer pairwise sequence similarities. This resulted in 122 pairwise similarity values for each read-to-read comparison. The average value for these 122 are shown in the cells of the grid (Fig. 3B). The highest within-read average was surprising low at 67% (for 782 × 782). In contrast, the between-read averages were 46% (for 782 × 801, 677, or 440).

Given that the sequences of the monomeric repeating units tended to vary within and between individual reads, we decided to use consensus sequence data to guide oligo FISH design (Fig. 3C). For high priority reads, we used the Tandem Repeats Finder program (Benson, 1999) to define read-specific consensus sequences. We next carried out multiple sequence alignments of these consensus sequences to identify the most highly conserved sequence regions which were considered ideal for design and production of fluorescent oligonucleotide probes (Fig. 3C). A list of new and previously published tandem repeats and FISH probes for hop are summarized in Table 2. Collectively, these represent the beginning of a new toolkit for hop cytogenomics, suitable for future investigations for structural genomics, segregation patterns, and chromosome evolution in hop. Their utility is demonstrated below using two of these new reagents, the oligo FISH probes for HuluTR120 and HuluTR225, in wild collected var. neomexicanus hop.

### Aberrant Meiosis And HuluTR FISH in Wild Hop

An important question in hop genome evolution is whether or not aberrant meiosis is a natural, intrinsic feature of hop or whether it can be explained entirely as a result of breeding and cultivation with structurally diverse genomes. To begin to address this issue and to demonstrate a possible application of these new FISH probes, wild hop was collected from what are thought to be isolated populations (Reeves and Richards, 2011) in the Arizona Sky Islands and male meiosis analyzed cytologically as shown in Figures 4–6.

**Figure 4.**
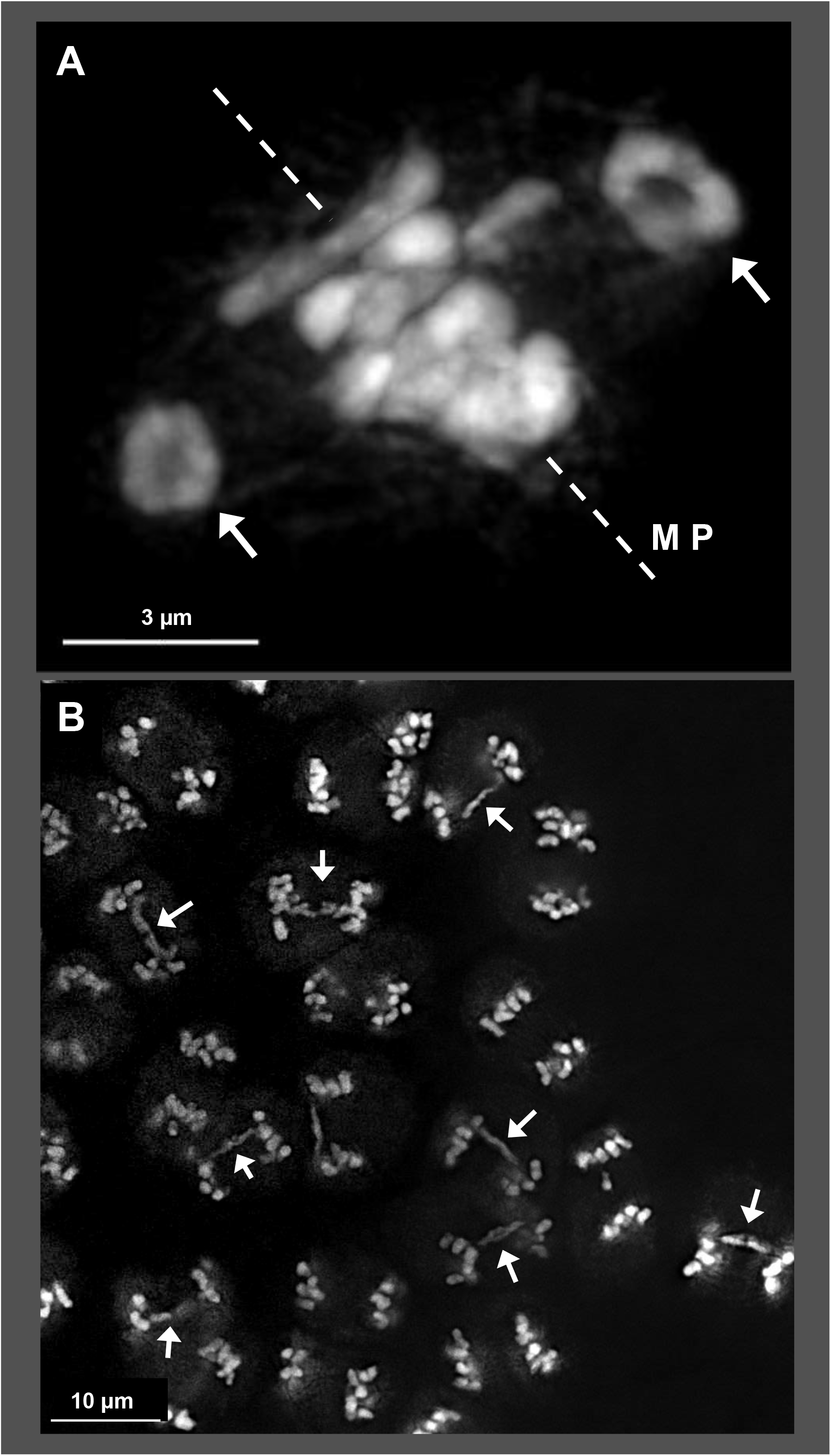
Meiotic abnormalities in Arizona Sky Island wild neomexicaus hop, plant TM2-82C. DAPI stained through-focus projections of (A) metaphase I, with bivalents (arrows) outside of the metaphase plate (MP), indicated by dashed line; (B) group of meiotic cells at anaphase I where half of the dividing nuclei exhibit anaphase bridges (arrows). The length of the scale bars are indicated in micrometers. More than 30 nuclei from plant TM2-82C were imaged and analyzed over multiple slides (n=6) during metaphase I. More than 30 nuclei from plant TM2-82C were imaged and analyzed over multiple slides (n=6) during anaphase I. Image in panel B was processed to correct for non-uniform staining (using Lightroom software).

We found evidence of aberrant chromosomal behavior at metaphase I (Fig. 4A) and anaphase I (Fig. 4B) using 3D imaging of DAPI-stained meiocytes. At metaphase, chromosomes typically congress on the metaphase plate, but in the example shown, two presumed ring bivalents (arrows, Figure 4A) are seen to be excluded from the metaphase plate, indicative of a chromosomal positioning problem. The frequency of irregularities wasconspicuous and occasionally extreme as seen in the low magnification image of 22 anaphase-stage cells from a single plant (TM2-82C from Mt. Bigelow), 11 of which exhibited chromosome bridges (arrows, Fig. 4B). Another way to track chromosome bivalency is through observation of nucleoli. When homologous NORs pair and synapse, their associated nucleoli fuse into a single nucleolus. Given that reported hop karyotypes have a single NOR locus (Karlov et al., 2003; Divashuk et al., 2014), we would expect that normal pairing would result in fusion of the homologous NORs to give one large nucleolar region by mid-prophase. However, tracking nucleoli number in mid-late meiotic prophase, we found that the meiocytes from wild hop (plant SH2 from Mt. Lemmon) could show either of two different patterns, single nucleoli (‘“n” in Fig. 5A) or double (“n1”, “n2” in Fig. 5B). Interestingly, the double nuclei occurred at an unusually high frequency, observed in 11 of the 22 nuclei imaged in 3D. We interpret the presence of double nucleoli as deviation from normal disomic, homologous pairing at the NOR regions. Consistent with this interpretation, we observed that the average nucleolar volumes were 14 μ^3^ for single-nucleolus cells (n=17 nucleoli) and 5 μ^3^ for double-nucleolus cells (n=16 nucleoli in 8 cells), a 2.8 fold difference.

**Figure 5.**
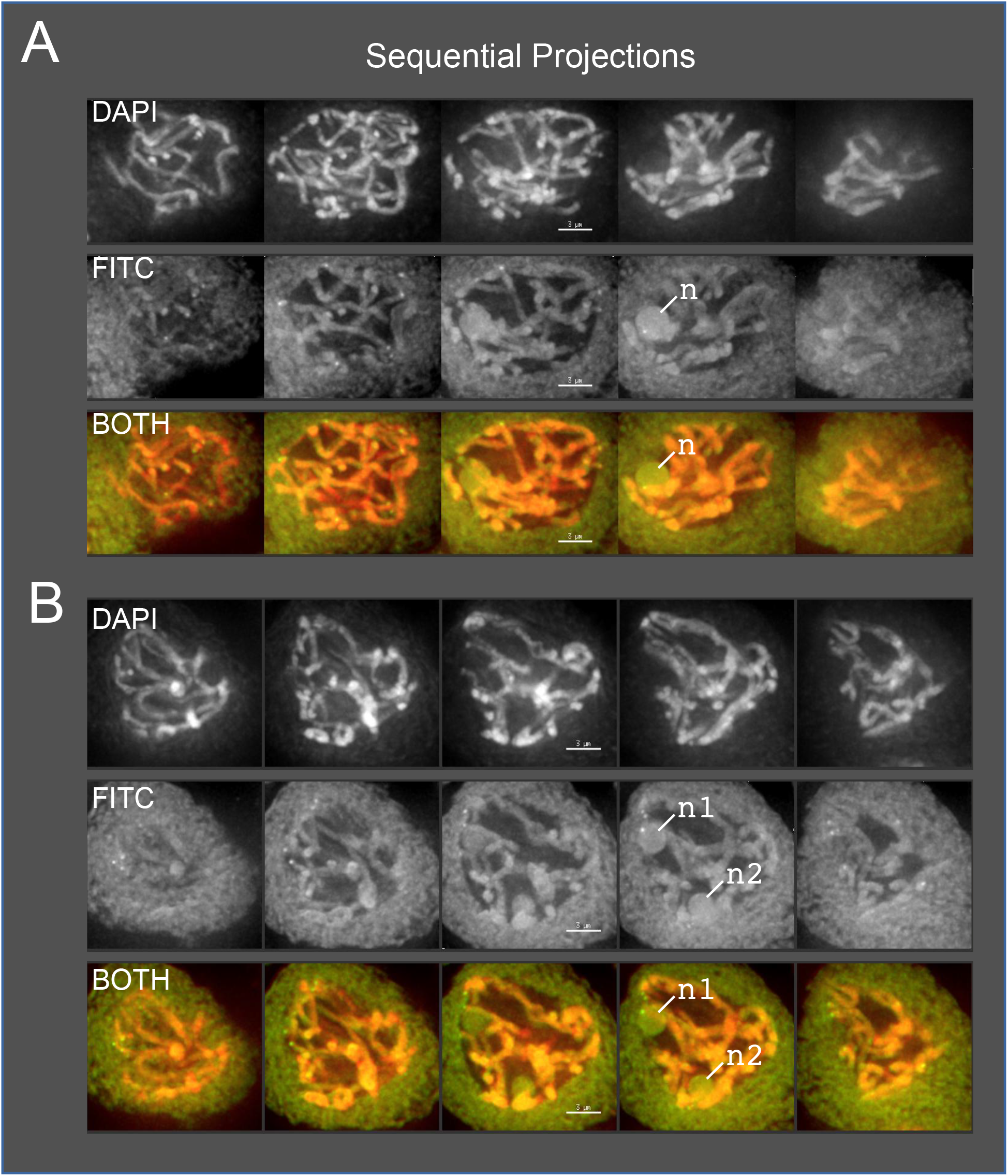
Single and double nucleoli during pachytene in Arizona Sky Island wild neomexicanus hop, plant SH2. Male flower buds were harvested and fixed in Farmer’s Fluid, then exchanged into Buffer A and formaldehyde fixed before microdissecting pollen mother cells from anthers for 3D acrylamide telomere FISH. The background fluorescence in the FISH channel reveals the location and number of nucleoli. Through-focus maximum-intensity sequential projections through two individual nuclei are shown in gray-scale for individual wavelengths or in color for overlay images, as labeled on the left. (A) Hop nucleus at mid-prophase showing a single nucleolus (‘n’ in FITC and BOTH). (B) Hop nucleus at mid-prophase showing two separate nucleoli (‘n1’, ‘n2’ in FITC and BOTH). The lengths of the scale bars (3 microns) are indicated. More than 70 nuclei from plant SH2 were imaged and analyzed over multiple slides (n=14) during pachytene.

**Figure 6.**
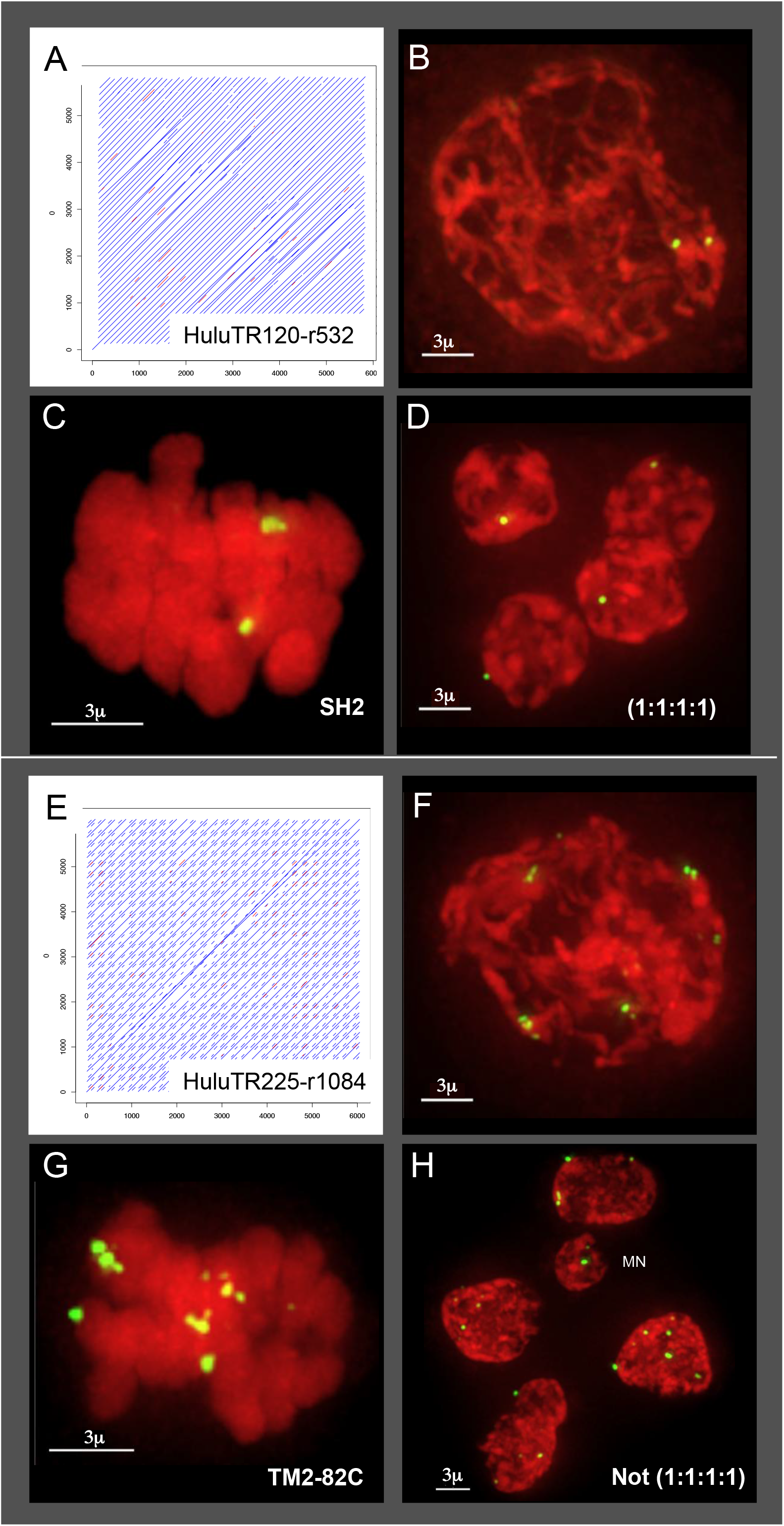
HuluTR120, 225 dot plots and FISH signals during meiosis in plants SH2 and TM2-82C. (A) Dot plot for HuluTR120. (B-D) Hop nucleus from plant SH2 hybridized with oligo FISH probe (TR120-Y) for HuluTR120 at (B) mid-prophase, (C) metaphase I and (D) tetrad stage. Meiotic prophase nuclei show two signals and an equal distribution of 1:1:1:1 after meiosis II at tetrad stage. (E) Dot plot for HuluTR225. (F-H) Hop nucleus from plant TM2-82C hybridized with oligo FISH probe (TR225-Y) for HuluTR225 at (F) mid-prophase, (G) metaphase I and (H) tetrad stage. Meiotic prophase nuclei show approximately 10-12 FISH signals per nucleus with variable size and brightness per signal spot. The tetrad-stage cell shows highly variable signals at the second meiotic division of approximately (5:4:2:4:7) and a micronucleus, labeled MN. The lengths of the scale bars are indicated in micrometers. More than 80 nuclei from plant SH2 were imaged and analyzed over multiple slides (n=5) stained with HuluTR120 during various meiotic stages. More than 80 nuclei from plant TM2-82C were imaged and analyzed over multiple slides (n=4) stained with HuluTR225 during various meiotic stages.

In order to test our new FISH probes on wild and non-Apollo hops, we applied two of them, HuluTR120 and HuluTR225, to meiocytes of two var. neomexicanus plants, SH2 and TM2-82C, as shown in Figure 6. We show that both of these HuluTR probes, designed from Apollo sequence data, successfully hybridized as discrete foci on the chromosomes of wild hop. In one case, the HuluTR120 probe gave two bright signals in plant SH2 as seen at mid-prophase (Fig. 6B) and metaphase I (Fig. 6C), a pattern indicative of paired homologous loci. At the tetrad stage, the HuluTR120 signals were distributed equally (Fig. 6D, 1:1:1:1). In another case, the HuluTR225 probe gave more complex patterns in plant TM2-82C, with variable brightness and size. The 10-12 FISH signals are seen at mid-prophase (Fig. 6F) and at metaphase I (Fig. 6G). The FISH signals appear to be distributed in an irregular pattern at both metaphase I and the post-meiotic tetrad-like stage (Fig. 6H). The examples represent multiple occurences of meiotic abnormalities from a single plant (Fig. 4B and Fig. 6E-H). Therefore, TR probes designed from one genotype can be used in others, and wild hops show both balanced (D) and unbalanced FISH signal distribution (H), similar to recent observations with 5S rDNA FISH (Easterling et al., 2018). his approach to develop new cytogenomic tools enabled the discovery and characterization of a class of tandem repeats with demonstrated utility for investigating the mysterious mechanisms of hop genome transmission and chromosomal evolution.

## DISCUSSION

Interest in tandem repeats has prompted investigators to develop new software programs to find or characterize tandem repeats using DNA sequencing data (Glunčić and Paar, 2013; Weiss-Schneeweiss et al., 2015; Mlinarec et al., 2019). Among the programs used are Tandem Repeats Finder (Benson, 1999), which uses string matching algorithms, and those utilizing graph-based clustering, such as RepeatExplorer (Novák et al., 2010) and TAREAN (Novák et al., 2017). These programs allow for the mining of existing and public repositories of genomic data to identify tandem repeats for various studies related to phylogenetics, genome evolution, and cytogenetics (Dodsworth et al., 2015; Belyayev et al., 2018; Mlinarec et al., 2019). More recently, long-read sequence data has been used to support FISH probe development in plants, with the aid of RepeatExplorer and TAREAN (Kapustová et al., 2019; Vondrak et al., 2019).

Here, we describe an approach using long-read sequences that allows for TR discovery aided by direct visual inspection of single self-aligned read dot plots. Even with these error-prone early generation single-molecule reads, we were able to uniquely and unambiguously find and group tandemly repeated sequence families and build consensus sequences. The DNA sequences from these reads were screened by k-mer analysis using criteria that yielded ~1000X enrichment for reads with the desired sequence features. The k-mer filtered dot plots provide highly informative way to visualize the data, making it easy to quickly interpret tandem repeat patterns within their genomic context one read at a time without any requirement for assembly. The data processing pipeline produced a PDF booklet of 1 read per page (Supplementary Material 1), which proved useful for downstream analysis. Compared to other methods, the approach described here has several notable advantages including (1) intuitive visualization of the genomic structure of the repeats, (2) highly sensitive ability to detect tandem repeats, as illustrated by the discovery of reads with HuluTR families present once per million reads (e.g. HuluTR050, HuluTR055, HuluTR070, HuluTR150, HuluTR280, and HuluTR350), (3) the retention of adjacent flanking genomic sequence, ideal for guiding genome assembly efforts, and (4) the retention of the individuality of TR clusters, which may come from multiple different loci. This last advantage may be helpful for future consideration of homologous alleles, homeologous alleles from hybrids, or multi-chromosomal loci on different paths of divergence. In contrast, the approach reported here has disadvantages such as the requirement for long-read sequences as the input data and the fact that the larger repeats, the less likely they will meet our k-mer threshold for 5-10 kbp reads. Given that we readily found known TRs (5S rDNA, HSR1, and telomere repeats) while also discovering 18 TRs (Table 1), on the whole, we consider this a robust approach to be widely applicable and relatively simple to interpret.

Repetitive sequences pose the greatest challenge for assembling complete genomes. The 1C genome size estimates for hop range from 2.5 - 3.0 Gb according to flow cytometric methods (Zonneveld et al., 2005; Grabowska-Joachimiak et al., 2006; Natsume et al., 2015) but only 2.1 Gb according to a recent from genome assembly (Natsume et al., 2015). Therefore, sequence assemblies currently account for only 80% of the known genome size, indicating that a large fraction of the genome is not represented in contemporary assemblies. Tandem repeat sequences are often mis-assembled and under-represented, being particularly prone to the repeat collapse problem in genome assembly. These discrepancies contribute to the genome size under-estimations while exacerbating problems associated with accurate contig assembly. For instance, markers flanking a TR cluster may be separated by only a few Kbp of TR, but reside on different contigs if only short read sequences guide the assemblies. Accurate incorporation of TR clusters is especially important in hop given its high degree of structural variability and segregation distortion (Zhang et al., 2017; Easterling et al., 2018).

Another useful aspect of defining TR families within long-read sequences is that they provide opportunities to explore the flanking genomic sequences for markers or genes. Sequences with “Partial read TR” or “Interspersed TR” patterns (Fig. 1) can be used to obtain non-TR regions for use in BLAST queries of genome databases. For instance, regions of non-TR sequences within the read housing HuluTR135 align with an EST of unknown function (GenBank Acc. FG346016.1) and map to a region (004949F:218,515․.248,514) in HopBase (Hill et al., 2017). Such approaches represent examples of new avenues to explore regions of biological importance for hop, such as sex chromosomes, segregation distortion hot spots, and key genes for flavor and aroma biosyntetic pathways or disease resistance.

A primary goal of this study was to produce new molecular cytology tools for hop chromosome research. To that end, we have described 18 new tandem repeat families (Table 1) and shown FISH results with probes for HuluTR120 and HuluTR225. To date, most of the hop chromosomes are numbered and distinguished by their relative size and in some cases their centromere locations as inferred from the primary constriction on mitotic chromosomes (Shephard et al., 2000). The most current hop karyotype includes HSR1, 5S rDNA, NOR, and telomere signals, which together uniquely tag 4 of the 10 chromosomes (Karlov et al., 2003; Divashuk et al., 2011, 2014). Notably, centromere-specific sequences have yet to be identified in hop, including this study. It is possible that among our HuluTR families are one or more that reside at centromeres. Alternatively, hop centromere repeats may not be organized as tandem repeats or their size and copy number may have resulted in their exclusion from our k-mer filtered subset of 1,121 reads. Indeed, a recent study in wheat found that centromeric tandem repeats enriched at CENH3 ChIP seq peaks can exceed 500 bp in repeat unit length (Su et al., 2019).

FISH probes are also invaluable for tracking meiotic chromosome interactions and post-meiotic transmission of discrete genetic loci. For instance, hop 5S rDNA FISH probes were previously used to document abnormal chromosomal interactions during pairing at late prophase and cytological segregation distortion in tetrads (Easterling et al., 2018). Here we present two new FISH probes that hybridize to a small number of discrete foci in wild hop plants. HuluTR120 FISH signals showed equal distribution of signals at the tetrad stage (1:1:1:1) in meiocytes from one wild plant, SH2. HuluTR225 FISH signals showed clear irregularities in meiocytes from a different wild plant, TM2-82C. An emerging picture is that there is considerable variation in FISH patterns even when using the same probe on cells from the one plant, siblings, or different varieties. This highlights the magnitude of the challenge of sorting out the hop genome and the importance of developing new markers of all types. With advances in hop genomics, and as the connections between physical chromosomes and linkage groups are elucidated, a cytological toolkit of TR FISH probes will accelerate an integrated view of the hop genome.

Wild hop populations occur naturally across the US in three varieties and are morphologically distinct but are not necessarily reproductively isolated (Reeves and Richards, 2011). They have been described as monophyletic (Tembrock et al., 2016) and are known to exhibit high levels of genetic diversity, particularly var. neomexicanus (Murakami et al., 2006). It is worth noting that cultivated, escaped hop plants, also referred to as ferals, can be mistaken for wild varieties, especially near areas where hop is cultivated or bred. In this study, we intentionally wanted wild neomexicanus hops and collected, therefore, from remote southwest US regions in the Arizona Sky Islands where the hop plants are morphologically distinct var. neomexicanus. Our cytological data in these wild plants (Figs. 4–6), together with previously reported meiotic segregation irregularities (Zhang et al., 2017; Easterling et al., 2018) establish that such meiotic abnormalities are clearly not limited to cultivated hop and can also occur in the wild. These findings, while limited in scope, highlight the recurrent observations of genomic instability in some members of the species. Similar phenomena have been observed Oenothera sp. and Clarkia sp., members of the Onagraceae family (Bloom, 1974; Hollister et al., 2019). Interestingly, some of these have stabilized structural variation though specialized meiotic behaviour possibly contributing directly to speciation events (Holsinger and Ellstrand, 1984). It remains to be determined whether the evolutionary dynamics of hop has contributed to speciation or divergence in the wild, questions that can be addressed using chromosome-marking FISH probes.

Here we increased the number of known tandem repeat sequence families in hop by nearly 10 fold using an innovative bioinformatic pipeline for de novo identification, visualization, and classification of TRs from long-read sequence data. This approach and the resulting cytogenetic resources should prove useful for further investigations into evolutionary, cytogenetic, or structural genomic research in hop.

## ACKNOWLEDGEMENTS

We thank Daniel Vera for help with the tandem repeat analysis. This work was supported by a Hopsteiner Doctoral Research Fellowship to KAE (FSU OMNI Award ID: 0000030675) and an FSU Planning Grant to HWB (FSU OMNI Award ID: 0000032134).

